# PWO1 interacts with PcG proteins and histones to regulate Arabidopsis flowering and development

**DOI:** 10.1101/226183

**Authors:** Mareike L. Hohenstatt, Pawel Mikulski, Olga Komarynets, Constanze Klose, Ina Kycia, Albert Jeltsch, Sara Farrona, Daniel Schubert

**Author notes:** These authors contributed equally (Mareike L. Hohenstatt, Pawel Mikulski). These authors contributed equally (Sara Farrona, Daniel Schubert). The author(s) responsible for distribution of materials integral to the findings presented in this article in accordance with the policy described in the Instructions for Authors (www.plantcell.org) are: Sara Farrona and Daniel Schubert.

## Abstract

Polycomb-group (PcG) proteins mediate epigenetic gene regulation by setting H3K27me3 via Polycomb Repressive Complex 2 (PRC2). In plants, it is largely unclear how PcG proteins are recruited to their target genes.

Here, we identified the PWWP-DOMAIN INTERACTOR OF POLYCOMBS1 (PWO1) protein which interacts with all three *Arabidopsis* PRC2 histone methyltransferases and is required for keeping full H3 occupancy at several Arabidopsis genes. PWO1 localizes and recruits CLF to nuclear speckles in tobacco nuclei, suggesting a role in spatial organization of PcG regulation. *PWO1* belongs to a gene family with three members acting redundantly: *pwo1 pwo2 pwo3* triple mutants are seedling lethal and show shoot and root meristem arrest, while *pwo1* single mutants are early flowering. Interestingly, PWO1’s PWWP domain confers binding to histones, which is reduced by a point mutation in a highly conserved residue of this domain and blocked by phosphorylation of H3S28. PWO1 carrying this mutation is not able to fully complement the *pwo1 pwo2 pwo3* triple mutant, indicating the requirement of this domain for PWO1 *in vivo* activity. Thus, the PWO family may present a novel class of histone readers which are involved in recruiting PcG proteins to subnuclear domains and in promoting Arabidopsis development.

## Introduction

Polycomb group (PcG) and the antagonistically acting Trithorax group (TrxG) proteins are key regulators of epigenetic gene regulation which are essential for the development of eukaryotic organisms (Kondo et al., 2016; Mozgova and Hennig, 2015). Initially identified in *Drosophila melanogaster,* PcG proteins maintain repression of homeotic gene expression, while the TrxG proteins sustain activation of homeotic genes through cell division, thus conferring epigenetic memory.

PcG proteins act in several high molecular weight complexes, the so-called Polycomb repressive complexes (PRC) (Mozgova and Hennig, 2015; Schuettengruber et al., 2007; Simon and Kingston, 2013). PRC2 consists of four core members in *Drosophila,* Enhancer of zeste (E(z)), Extra sex combs (Esc), p55 and Suppressor of zeste12 (Su(z)12). The PRC2 complex mediates tri-methylation of H3K27 through the SET domain of its subunit E(z) (Cao et al., 2002; Cao and Zhang, 2004; Czermin et al., 2002; Kuzmichev et al., 2002; Muller et al., 2002). Although the H3K27me3 mark is required for gene repression, the presence of this mark does not always correlate with the transcriptional status of the gene where it is present indicating a more complex regulation and the involvement of other factors (Bouyer et al., 2011; Farrona et al., 2011; Lafos et al., 2011). An additional complex, PRC1, has been also related to this activity. The PRC1 subunit Polycomb (Pc) specifically binds to H3K27me3 via its chromodomain. Thus, in the classical PcG model, PRC1 recognizes the presence of this histone mark and then inhibits nucleosome remodeling and transcription, compacts chromatin and ubiquitinates histone H2A (de Napoles et al., 2004; Fischle et al., 2003; Francis et al., 2004; Francis et al., 2001; Shao et al., 1999; Wang et al., 2004). However, more recent data indicate that the hierarchical recruitment of PRCs may be far more complex. Indeed, studies in plants and animals showed that PRC1 activity is required for proper H3K27me3 deposition to specific targets and PRC1 components have been found directly interacting with PRC2 components (Del Prete et al., 2015; Merini and Calonje, 2015; Schwartz and Pirrotta, 2014). In addition, it has been recently shown that PRC1-mediated repression can also occur in the absence of H2A monoubiquitination indicating different PcG silencing scenarios (Calonje, 2014; Illingworth et al., 2015; Pengelly et al., 2015).

Recruitment of PcG proteins to the chromatin may occur by several mechanisms, including binding of H3K36me3 by the PRC2 associated protein Polycomb-like, binding to transcription factors and to CpG dinucleotides in CpG islands, interaction of PhoRC with specific DNA sequences, the Polycomb response elements (PREs), and direct binding of long noncoding RNAs by PRC2 (Cai et al., 2013; Del Prete et al., 2015; Deng et al., 2013; Klose et al., 2013; Schuettengruber and Cavalli, 2009; Simon and Kingston, 2013). PRE-like sequences have been identified at a few loci in plants and the presence of specific cis-regulatory elements (e.g. GAGA motives, telo boxes) in these sequences have been related with PcG recruitment (Berger et al., 2011; Deng et al., 2013; Hecker et al., 2015; Lodha et al., 2013; Wang et al., 2016; Zhou et al., 2016). Binding of PRC2 and PRC1 components to long noncoding RNAs has also been shown in plants (Ariel et al., 2014; Heo and Sung, 2011). Nevertheless, it remains elusive whether these recruitment mechanisms generally apply for the thousands of PcG target genes in Arabidopsis.

All four subunits of the *Drosophila* PRC2 complex are conserved in Arabidopsis. Except for the single copy gene *FERTILIZATION INDEPENDENT ENDOSPERM (FIE)* which is the ortholog of Esc, all subunits are encoded by small gene families (Mozgova and Hennig, 2015; Ohad et al., 1999). The catalytic subunit E(z) is represented by three genes in the Arabidopsis genome encoded by *CURLY LEAF (CLF), SWINGER (SWN)* and *MEDEA (MEA)* (Goodrich *et al.,* 1997;Grossniklaus *et al.,* 1998;Luo *et al.,* 1999). In addition, Su(z)12 is orthologous to the Arabidopsis genes *VERNALIZATION 2 (VRN2), EMBRYONIC FLOWER 2 (EMF2)* and *FERTILIZATION INDEPENDENT SEED 2 (FIS2)* (Gendall et al., 2001; Luo et al., 1999; Yoshida et al., 2001). The genes *MULTICOPY SUPPRESSOR OF IRA 1-5 (MSI1-5)* have sequence homology to the WD40 protein p55, and MSI1 and MSI4 were found to be associated with *Arabidopsis* PRC2 complex (Derkacheva et al., 2013; Kohler et al., 2003; Pazhouhandeh et al., 2011). Genetic studies suggest the existence of at least three PRC2 complexes in *Arabidopsis:* the VRN-PRC2 complex which is involved in the vernalization (vrn) response, the FIS-PRC2 complex required to inhibit fertilization independent seed development (fis) and the EMF-PRC2 complex that is needed for the suppression of precocious, embryonic flowering (emf) and for floral organ development (Forderer et al., 2016; Mozgova and Hennig, 2015). Hence the different complexes control important transitions at different stages of plant development and are, therefore, crucial to accomplish the complete life cycle (Butenko and Ohad, 2011; Forderer et al., 2016; Mozgova et al., 2015). Genome-wide profiling of PRC2 target genes has revealed the existence of more than 8000 genes carrying the H3K27me3 mark in *Arabidopsis* (Bouyer et al., 2011; Lafos et al., 2011; Oh et al., 2008; Roudier et al., 2011; Zhang et al., 2007). These studies identified previously known target genes of the three potential PRC2 complexes, however, also identified targets involved in stress responses, in hormonal signal transduction pathways, in promotion of embryonic growth and microRNA genes, suggesting that additional PRC2 complexes may exist.

It is likely that plant-specific components of PRCs have evolved to fulfill similar function to the non-conserved Drosophila PcG proteins or functions specific to plant growth and development. Recently, several PRC2 components have been identified which modulate PRC2 activity: VRN5 and VIN3 associate with PRC2 to form a PHD-PRC2 to achieve high levels of H3K27me3, similar to Drosophila Pcl-PRC2 and human hPHF1-PRC2 (Greb et al., 2007; Nekrasov et al.; 2007; Cao et al.; 2008; Sarma et al.; 2008). Additional PcG-associated proteins encoded only in plant genomes have been either identified in genetic screens for suppressors of *lhp1* and *clf* (ANTAGONISTIC OF LHP1 (ALP1); TELOMERE REPEAT BINDING (TRB) and an Arabidopsis *Chromosome transmission fidelity 4 (Ctf4)* homologue) and by protein-protein interactions screens (BLISTER (BLI), ALFIN1-like proteins (ALs)) (Liang et al., 2015; Molitor et al., 2014; Schatlowski et al., 2010; Sung and Amasino, 2004; Zhou et al., 2016; Zhou et al. 2017). Although ALP1 is found in PRC2 complexes, it antagonizes PRC2 function, suggesting that PRC2 activity can be modulated at various levels. However, how the other PRC2 associated proteins molecularly contribute to PcG silencing is largely unresolved.

Thus, PcG target genes are regulated at multiple levels including recruitment of PcG proteins, a combination of multiple repressive modifications, absence of activating modifications and binding/interpretation of the marks. While the role of PRC2 and PRC1-like proteins in plant development is relatively well understood now, many regulators controlling additional molecular functions in PcG silencing are awaiting their discovery.

In this study, we present the identification of the novel, plant-specific PWWP domain protein PWWP-DOMAIN INTERACTOR OF POLYCOMBS (PWO1), which interacts with all three PRC2 histone methyltransferase subunits from *Arabidopsis.* PWO1 localizes to euchromatic regions in tobacco nuclei, both in the nucleoplasm and in nuclear speckles. PWO1 contributes to PcG silencing by repressing a subset of PcG targets. While H3K27me3 levels are reduced at these loci in *pwo1* mutants, this is largely explained by a reduction in H3 occupancy, suggesting that PWO1 contributes to chromatin compaction. *pwo1* mutants are early flowering due to reduced levels of the floral repressor *FLOWERING LOCUS C (FLC).* We show that PWO1 has a much broader role in development as it acts redundantly with two homologous proteins to maintain shoot and root meristems. Interestingly, the putative chromatin reading PWWP domain of PWO1 is required for nuclear speckle formation and confers binding to histone 3 (H3) *in vitro.* This function is inhibited by phosphorylation of H3S28, a mark counteracting PcG silencing. A point mutation of PWO1’s PWWP domain strongly reduces PWO1-H3 interaction *in vitro* and, when transformed in the *pwo1 pwo2 pwo3* background, is unable to fully complement the triple mutant inducing developmental abnormalities at the shoot apical meristem. Thus, we identify the PWO family as essential regulators of Arabidopsis development which may recruit PRC2 to specific regions on the chromatin by interacting with H3 through its PWWP domain.

## Results

### Identification of PWO1 as an interactor of PcG proteins

To discover proteins involved in PcG mediated gene silencing, we performed a yeast two-hybrid screen with a truncated CLF protein as bait (Schatlowski et al., 2010). This screen yielded the Su(z)12 homologs EMF2 and VRN2 and the PcG associated protein BLISTER (Schatlowski et al., 2010). We revisited the list of potential CLF interaction partners to identify proteins containing putative chromatin “reading” domains. This analysis identified the protein At3g03140 which contains 769 amino acids and comprises a predicted amino-terminal PWWP domain as well as a central nuclear localization signal (NLS) (Figure 1B). PWWP domains belong to the ‘Royal family’ which includes the Tudor, Chromatin-binding (Chromo), Malignant Brain Tumor (MBT), PWWP and Agenet domains and are implicated in methyl-lysine/arginine binding (Maurer-Stroh et al., 2003). Subsequently, we confirmed the interaction with CLF in independent yeast two-hybrid analyses and further revealed a potential interaction with the two homologs of CLF, SWINGER (SWN) and MEDEA (MEA) and homo-dimerization of At3g03140 (Figure 1A). We therefore named At3g03140 PWWP-DOMAIN INTERACTOR OF POLYCOMBS 1 (PWO1). To confirm the interaction *in vivo,* we generated *PWO1_pro_::PWO1-GFP*i*35S_pro_::mCHERRY-CLF* double transgenic Arabidopsis lines. The *i35S_pro_::mCHERRY-CLF* line allows estradiol-dependent induction of mCHERRY-CLF and, therefore, controlled expression levels. After estradiol induction, proteins were isolated and subjected to co-immunoprecipation analyses. Anti-GFP antibodies precipitated PWO1-GFP and also pulled down mCHERRY-CLF suggesting that PWO1 and CLF are part of the same complex *in planta* (Figure 1C).

**Figure 1:**
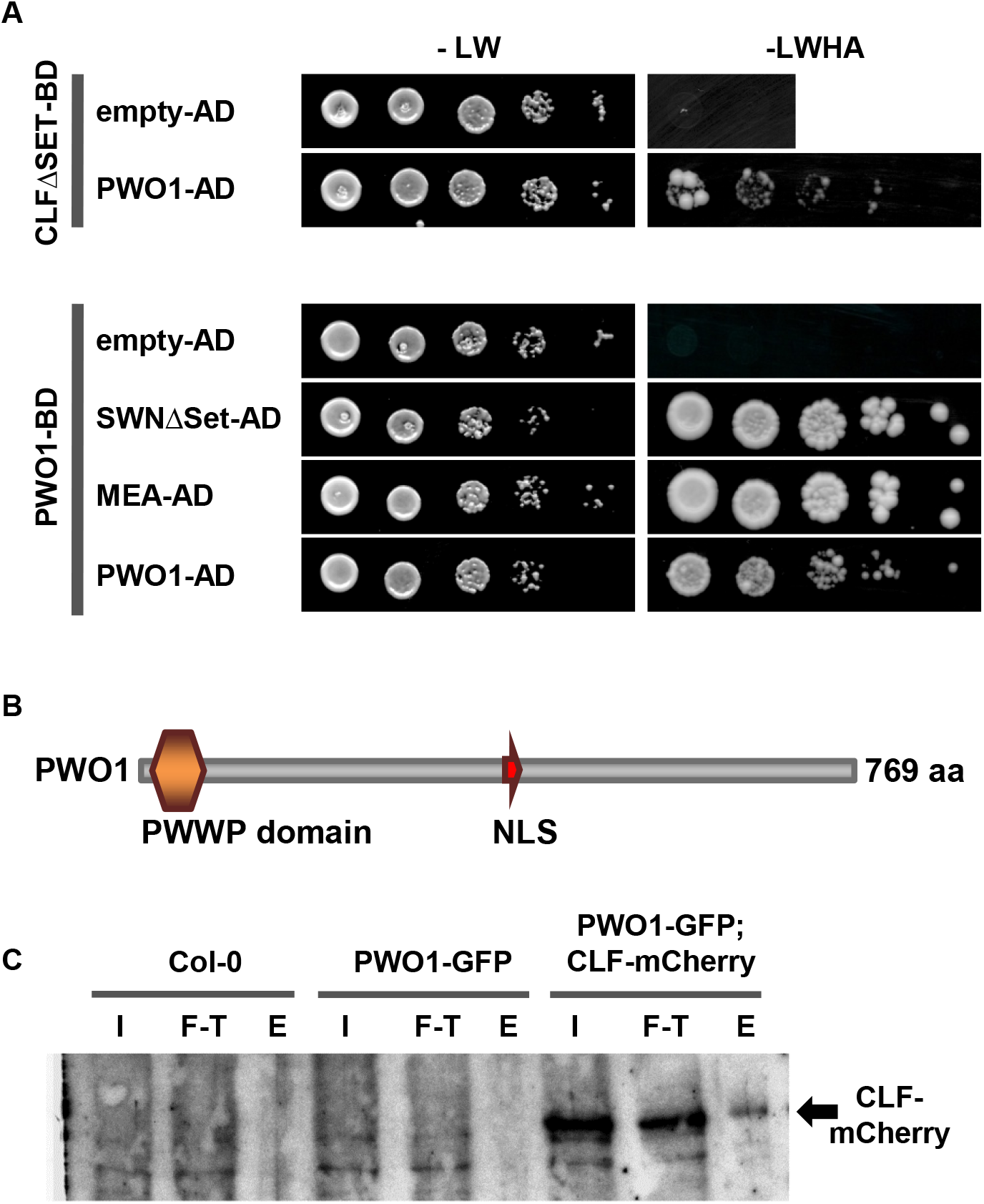
Interaction of PWO1 with PRC2 members. **A**. Yeast-two hybrid analyses detect an interaction of PWO1 with CLF, SWN and MEA and PWO1 homodimerization. Yeasts containing the various combination were grown on medium selecting for plasmids (-LW (-Leucine, Tryptophane) or for reporter gene activation (-LWAH; Adenine, Histidine). Serial dilutions are shown. BD: GAL4-DNA-binding fusion, AD: GAL4-DNA-activation domain fusion. For CLF and SWN, constructs lacking the SET domain were taken (ΔSET). **B**. Schematic presentation of the predicted PWO1 protein. PWWP: Proline-Tryptophane-Tryptophane-Proline domain; NLS: nuclear localization signal. **C**. Immunoblot derived from co-immunoprecipitation of PWO1 and CLF from stable Arabidopsis transgenic lines. IP was performed with anti-GFP antibodies, detection was with anti-mCherry antibodies. I: input, F-T: flow-through, E: eluate.

While PWWP-domain proteins are found in most eukaryotic organisms, proteins with high similarity to PWO1 are only found in plants including two close Arabidopsis homologs that we named *PWO2* (At1g51745) and *PWO3* (At3g21295) (supplemental Figure S1). An alignment of the protein sequences of PWO1/2/3 shows the highest similarity in the amino-terminal region of the proteins containing the predicted PWWP domain and in the C-terminal region (supplemental Figure S1).

### PWO1 is widely expressed and tethers CLF to nuclear speckles in tobacco

In order to analyze the expression pattern of *PWO1*, a translational fusion of the PWO1 genomic locus to a carboxy-terminal uidA reporter gene (*PWO1_pro_::PWO1-GUS*) was constructed and introduced in Col-0 plants by Agrobacterium mediated transformation. Analysis of homozygous *PWO1_pro_::PWO1-GUS* reporter lines revealed *PWO1* expression in the vasculature of the root and the shoot, cotyledons and rosette leaves of young seedlings. In addition, expression was observed in the primary root tip as well as in developing side roots. In flowers expression of the fusion protein was detectable in sepals and carpels (Figure 2B). Thus *PWO1* expression is widely expressed, preferentially in less differentiated tissues, and broadly overlaps with that of the PcG genes *CLF* and *SWN* (Chanvivattana et al., 2004; Goodrich et al., 1997).

**Figure 2:**
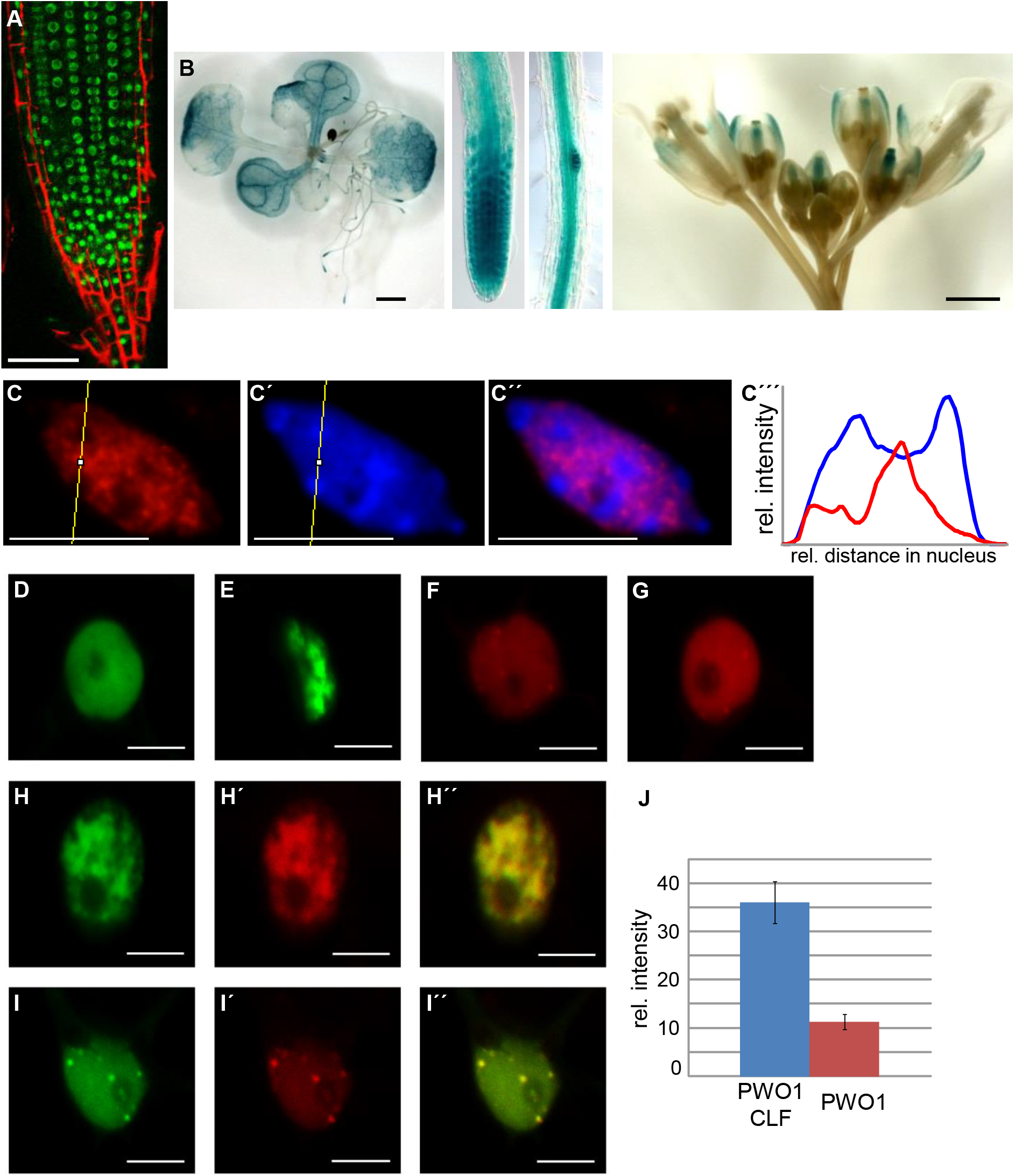
PWO1 localization, expression pattern and co-localization with CLF. **A**. *PWO1_pro_.:PWO1-GFP* reveals PWO1 nuclear localization and expression in most cells of the root tip. **B**. *PWO1_pro_::PWO1-GUS* line, GUS staining is detected in seedlings, root tips, root vasculature and inflorescences. **C**. Immunofluorescence of nucleus isolated from *PWO1_pro_::PWO1-GFP* seedlings, stained with anti-GFP (C) and DAPI (C’); C’’ is merge of C and C’; C’’’ depicts the profiles of anti-GFP (red) and DAPI (blue) fluorescence intensities through the yellow line in c and C’. **D. - I**. Transient expression in *N. benthamiana* leaf epidermal cells. Expression was induced with 2 μM β-estradiol for at least 5 hours. **D., E**. *i35S_pro_:GFP-CLF;* **F., G.** *i35S_pro_:PWO1-mCherry,* **H., I**. Co-expression of *i35S_pro_:GFP-CLF* and *i35S_pro_:PWO1-mCherry,* H’’ and I’’ are merges of the two channels. Scale bars: A. 50 μM, B. 1 mm, C. 5 μM, D-I 10 μM.. **J.** Intensity profiles of speckles in *i35S_pro_:PWO1-mCherry* tobacco nuclei (N=6) and *i35S_pro_:GFP-CLF,i35S_pro_:PWO1-mCherry* co-transformed nuclei (N=6).

As PcG proteins are localized to euchromatic regions of the nucleus, we asked whether PWO1 shows similar localization. We therefore analyzed PWO1 localization in a *PWO1_pro_.:PWO1-GFP* Arabidopsis line. Consistent with the predicted NLS, the PWO1-GFP fusion protein showed nuclear localization in *Arabidopsis* root cells (Figure 2A). In addition, immunofluorescence analyses on isolated nuclei of the *PWO1_pro_::PWO1-GFP* line revealed a non-uniform nuclear distribution of PWO1 and uncovered an exclusion from the heterochromatic chromocenters which are densely stained by the DNA stain DAPI (4’,6-Diamidine-2-phenylindole) suggesting that PWO1 is a euchromatic protein (Figure 2C).

In order to assess whether PWO1 co-localizes with its interaction partner CLF, PWO1 - mCherry (*i35S_pro_::PWO1-mCherry)* was transiently co-expressed with GFP-CLF (*i35S_pro_::GFP-CLF*) in *Nicotiana benthamiana.* Both fusion proteins localized to the nucleus which is consistent with the interaction of PWO1 with CLF (Figure 2D-I). Interestingly, PWO1-mCherry showed localization not only to the nucleoplasm but also to a variable amount of nuclear speckles of unknown identity. GFP-CLF localized either uniformly or in larger patches to the nucleus but never to speckles when expressed alone. However, when PWO1-mCherry was co-expressed with GFP-CLF, one third of nuclei (5 of 15 nuclei) showed speckle formation for GFP-CLF which are largely overlapping with the PWO1 speckles. In addition, another third of nuclei (5 of 15 nuclei) displayed localization of both PWO1 and CLF to larger patches in the nucleus which was not observed in PWO1-mCherry single transformations. Analyses of the confocal images demonstrated an increase in speckle intensity when both proteins were co-transformed together (Figure 2J). Hence, subnuclear localization of both PWO1 and CLF in a heterologous system depends on each other and provides further evidences that they may form a complex *in vivo.*

### Role of the PWO family in flowering time and seedling development

In order to analyze PWO1 function during *Arabidopsis* development, two T-DNA insertions in the *PWO1* gene and a line carrying a premature stop-codon in the PWWP domain were isolated and homozygous plants analyzed. As the stop-codon results in a truncated protein of only 45 amino acids, this allele likely reflects a loss of function allele (supplemental Figure S2). All three *pwo1* alleles had no obvious leaf or flower defects, however they consistently showed a moderate early flowering phenotype in long (LD) and short day (SD) conditions which could be complemented by introducing *PWO1_pro_::PWO1-GFP* (Figure 3A-C and supplemental Figure S3).

**Figure 3:**
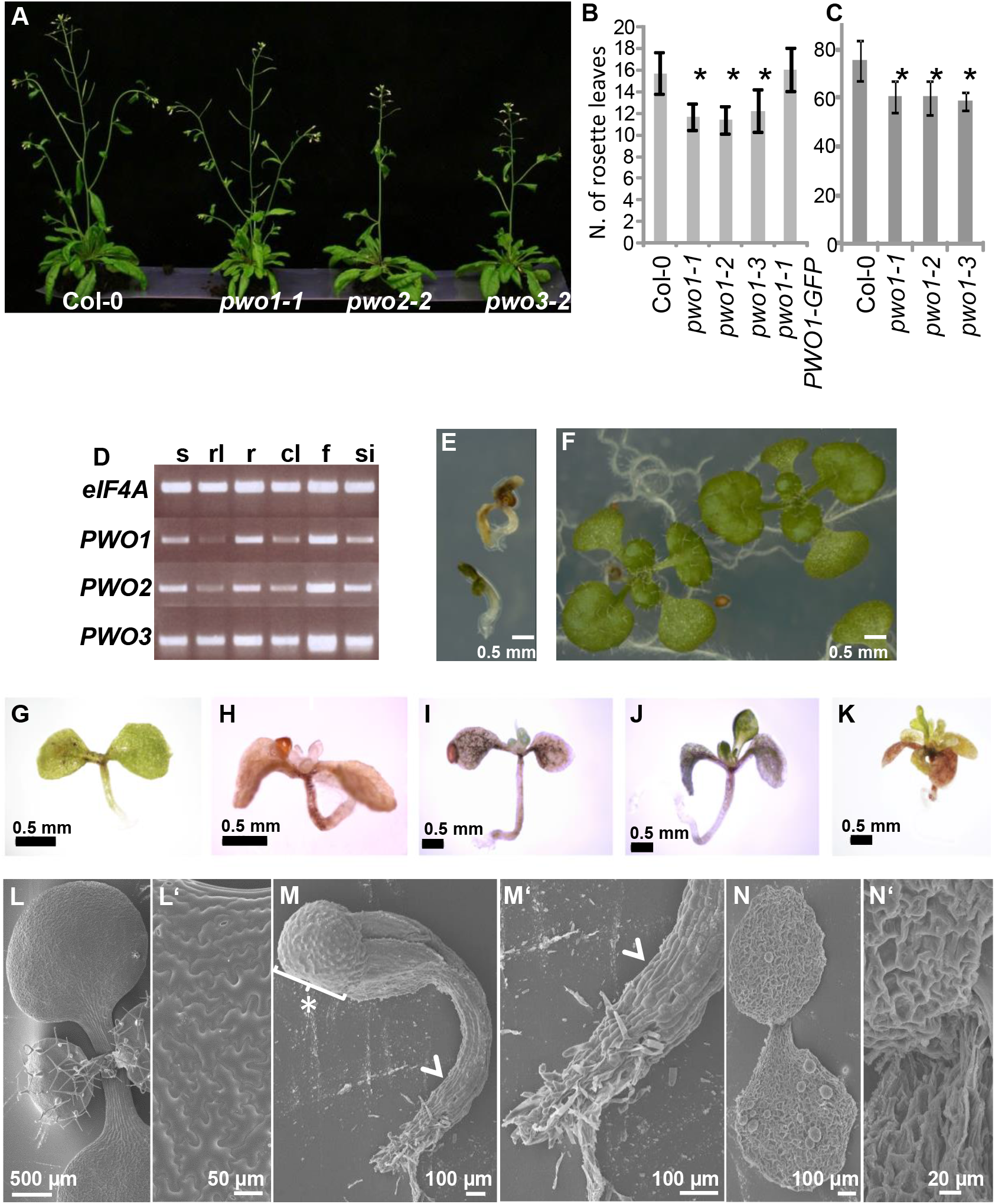
Developmental analyses of *pwo1, pwo2, pwo3* single and *pwo1/2/3* triple mutants. **A.** Growth phenotype of wildtype and *pwo1-1, pwo2-2, pwo3-2* plants (30 dag). **B, C.** Flowering time analyses of wildtype Col-0, *pwo1* alleles and *pwo1-1 PWO1_pro_::PWO1-GFP (pwo1-1* PWO1-GFP) in LD (B) and SD (C), x-axes indicate number of rosette leaves at time of bolting; n≥19; asterisks indicate significantly different number of rosette leaves compared to Col-0 (Student’s t-Test, p<0.01). **D.** Semi-quantitative expression analyses (RT-PCR) of *PWO1, PWO2* and *PWO3* in seedlings (s), rosette leaves (rl), roots (r), cauline leaves (cl), flowers (f) and siliques (si), *eIF4A* was used as reference gene. **E, F.** *pwo1/2/3* triple mutants show seedling lethality (E) which is rescued by a *PWO1_pro_::PWO1-GFP* transgene (F). Plants 14 dag are shown.. **G-K.** Various classes of seedling phenotypes of *pwo1/2/3* triple mutants (10 to 21 dag). **L** SEM analyses of wildtype seedling (14 dag), L’ close-up of cotyledon epidermis. **M**. *pwo1/2/3* seedling 14 dag, note non-collapsed epidermal cells (arrowhead); asterisk with brackets indicates seed coat; M’ close-up of root tip in H, **N**. *pwo1/2/3* seedling 21 dag, note collapsed epidermal cells, N’ close-up of SAM shows lack of leaf primordia.

The analysis of the morphological phenotypes of *pwo1* mutant alleles showed only mild phenotypes compared to PcG mutants like *clf* or *lhp1/tfl2* (Goodrich et al., 1997; Larsson et al., 1998). Since *PWO1* is a member of a gene family, the expression patterns of the two genes with closest similarity to *PWO1, PWO2* and *PWO3,* were analyzed by RT-PCR. RNA from different tissues was isolated, and all three genes showed a largely identical expression pattern in all tissues that were tested (Figure 3D). While T-DNA insertions in *PWO2* and *PWO3* and double mutant combinations *pwo1 pwo2, pwo1 pwo3* and *pwo2 pwo3* resulted in mild morphological phenotypes compared to the wildtype, *pwo1-1 pwo2-2 pwo3-2* triple mutants exhibited a dramatic seedling phenotype (Figure 3E and G-K and supplemental Figure S2). The triple mutants showed a termination of the apical root and shoot meristems soon after germination and accumulation of anthocyanins in the shoot tissues (Figure 3E and G-K). Most seedlings died 14 dag, lacked chlorophyll and appeared brownish or translucent. Analysis of the triple mutants by scanning electron microscopy (SEM) uncovered that epidermal cells of the triple mutants are collapsed around 14 dag. These cells appeared non-collapsed 7 dag, suggesting that this phenotype occurred post-embryonically (Figure 3L-N). *pwo1-1 pwo2-2 pwo3-2* triple mutants rarely produced leaf-like organs which appeared needle-like and usually lacked trichomes. To check whether this phenotype depends on *PWO* genes, we introduced *PWO1_pro_.:PWO1-GFP* or *PWO3_pro_::PWO3-GUS* transgenes in the triple mutants which both fully rescued the triple mutant phenotype (Figure 3F and supplemental Figure S2). Collectively, our analyses uncover an essential role for the *PWO* gene family in post-embryonic growth and in maintenance of both root and shoot meristems; however, this function is masked by the redundancy of *PWO1, PWO2* and *PWO3.*

### *PWO1* interacts genetically with *CLF* and regulates expression of PcG target genes

The physical interaction of PWO1 and CLF suggested that PWO1 and CLF have overlapping functions during *Arabidopsis* development. To test this, we generated *pwo1-1 clf-28* double mutants, which resulted in a strong enhancement of the *clf* phenotype: a severe reduction of plant size, very strong upwards leaf curling and day length independent early flowering suggesting a genetic interaction of *PWO1* and *CLF* (Figure 4A-C). *pwo1* mutants strongly enhanced the *clf* phenotype even in short day conditions where the *clf* leaf phenotype is largely suppressed (supplemental Figure S3).

**Figure 4:**
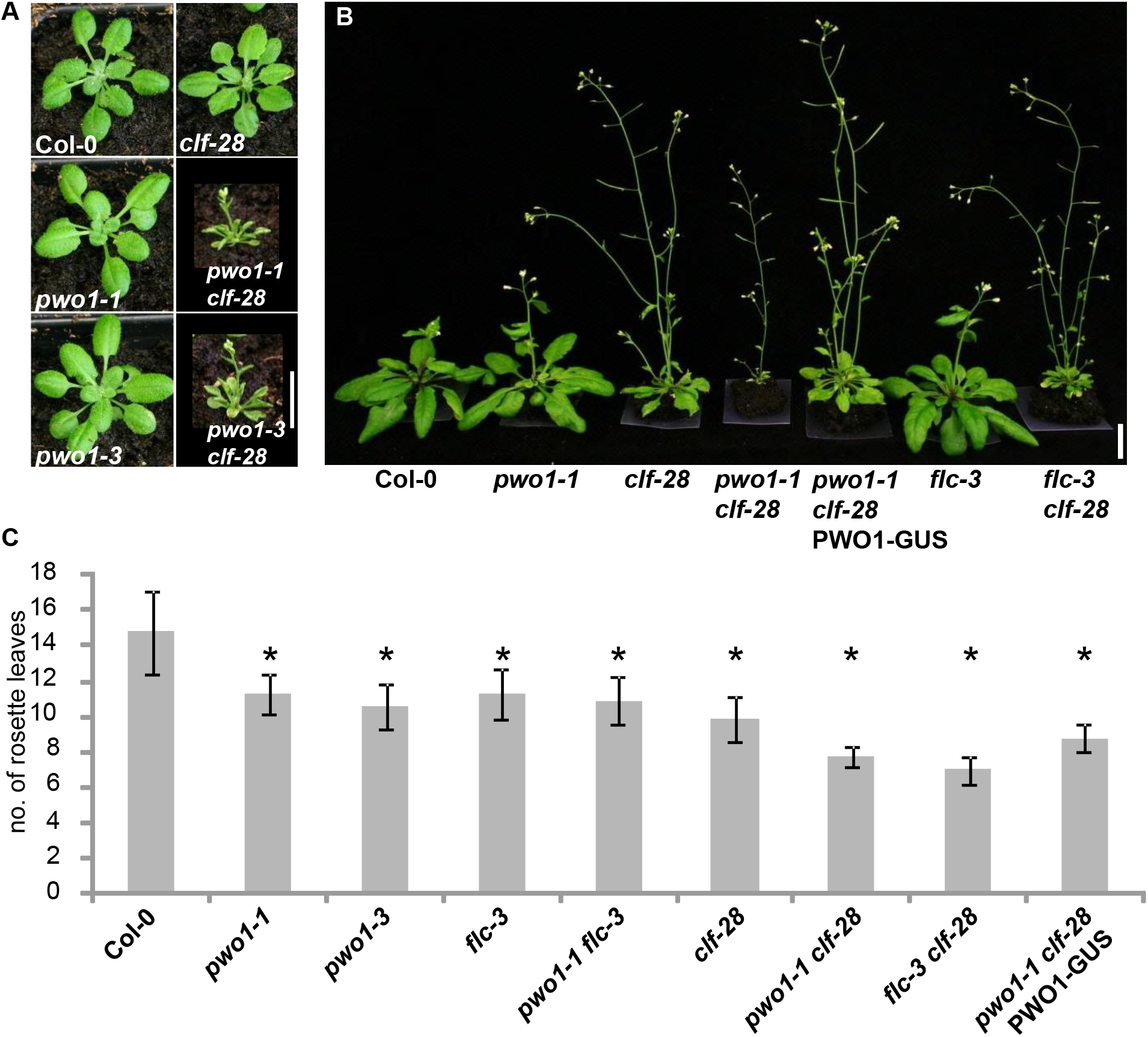
Genetic interaction of *pwo1, clf* and *flc* and regulation of PcG target genes by PWO1. **A.** *pwo1* alleles enhance the *clf* mutant phenotype (plants 21 dag, grown in LD conditions). **B.** *pwo1* and *flc-3* enhance the *clf-28* phenotype, but *pwo1-1 clf-28* double mutants have a stronger phenotype than *flc-3 clf-28* (plants 30 dag, LD conditions). **C.** Flowering time analyses of plants grown in LD, loss of *PWO1* and *FLC* similarly enhance early flowering of *clf-28,* while *pwo1-1, flc-3* and *pwo1-1 flc-3* are similarly early flowering. n≥23 (except for *clf-28 flc-3;* n=3); asterisks indicate significantly different number of rosette leaves compared to Col-0 (Student’s t-Test, p<0.001). Rosette leaf numbers of *pwo1-1, pwo1-3, flc-3* and *pwo1-1 flc-3* are not significantly different to each other (Student’s t-Test, p>0.1).

To reveal the genes causal for enhancement of the *clf* mutant phenotype we tested expression of the MADS box genes *FLOWERING LOCUS C (FLC), FLOWERING LOCUS T(FT), AGAMOUS (AG)* and *SEPALLATA3 (SEP3)* in *pwo1* single and *pwo1 clf* double mutants. Ectopic expression of *AG* and *SEP3* are largely responsible for the *clf* mutant phenotype, while loss of *FLC* enhances it (Goodrich et al., 1997; Jiang et al., 2008; Lopez-Vernaza et al., 2012). Consistent with day-length independent early flowering of *pwo1* and *flc* mutants (Figure 3B-C) (Michaels and Amasino, 1999; Sheldon et al., 1999), *FLC* was hardly detectable in *pwo1* (Figure 5A), whereas *FT* expression was not affected (Supplemental Figure 4). In addition, *pwo1 flc* double mutants showed similar flowering time as each single mutant suggesting that *flc* is epistatic to *pwo1* in LD and SD conditions in terms of flowering time control (Figure 4C and Supplemental Figure S3). As an introduction of *flc-3* in the *clf-28* mutant background leads to an enhancement of the early flowering and leaf curling phenotype (Lopez-Vernaza et al., 2012) the enhancement of *clf* by *pwo1* may be largely due to lower levels of *FLC.* Indeed, *pwo1-1 clf-28* mutants showed similar mis-regulation of *FT* and flowered at the same time as *flc-3 clf-28,* indicating that the decrease of *FLC* expression in the *pwo1-1* mutant might be responsible of enhancing the *clf-28* flowering phenotype (Figure 3 and Supplemental Figure 4). However, *pwo1-1 clf-28* mutants showed a stronger enhancement of the *clf-28* phenotype in respect to plant size and leaf curling than the *flc-3 clf-28* double mutants (Figure 4B), suggesting that mis-regulation of additional genes besides *FLC* is causal for enhancement of *clf-28* by *pwo1-1.* We therefore analyzed expression of the PcG target genes *AG* and *SEP3* whose mis-expression is largely responsible for the *clf* mutant phenotype (Goodrich et al., 1997; Lopez-Vernaza et al., 2012). Importantly, *AG* and *SEP3* showed a similar mis-expression in *flc-3 clf-28* compared to the *clf-28* single mutant, while in *pwo1-1 clf-28 AG* was slightly and *SEP3* was more strongly expressed than in *clf-28* (Figure 5A). To reveal whether changes in PcG target gene expression are correlated with reduced levels of the PRC2 mark H3K27me3, we performed chromatin immunoprecipitation (ChIP) experiments in Col-0, *pwo1* and *pwo1 pwo3.* While H3K27me3 occupancy was significantly reduced at *SEP3, AG* and *FUSCA3* in single and double PWO mutants compared to Col-0 (Figure 5B and supplemental figure S5), *FLC* H3K27me3 levels were only affected in *pwo1 pwo3* double but not in *pwo1.* However, we observed an even stronger reduction in H3 occupancy at all tested loci, suggesting that PWO1 is rather required for full nucleosomal levels than for high level H3K27me3. In summary, *SEP3* and *AG* upregulation and *FLC* downregulation in *pwo1-1 clf-28* double mutants compared to *clf-28* single mutants largely explain the double mutant phenotype. In addition, PWO1 represses *SEP3* at least partially independently of FLC and regulates H3K27me3/H3 enrichment at the tested loci

**Figure 5:**
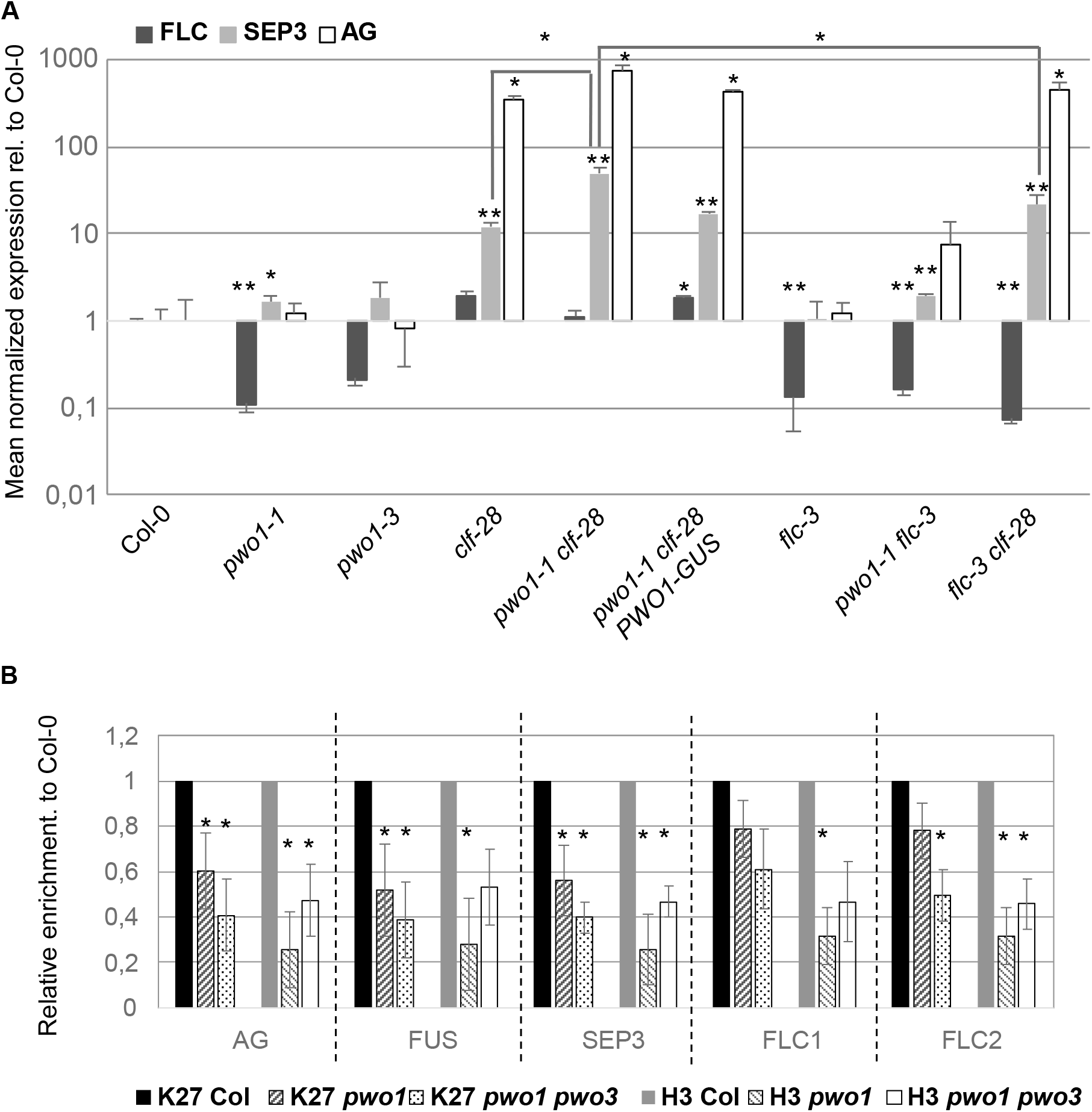
Expression and chromatin analyses in *pwo1* mutants. A, B. qRT-PCR analyses of the PcG target genes *FLC, SEP3* and *AG* in various mutant backgrounds. Pools of seedlings grown in LD were harvested 10 dag. Asterisks indicate significant difference (p<0.01 for *; p<0.001 for **, Student’s t-Test), significance was analysed in comparison to wild-type (Col-0) or as indicated by brackets; error bars: standard error of three biological replicates, grown and harvested independently. Note logarithmic scaling. C. Chromatin Immunoprecipitation in wild-type, *pwo1* and *pwo1 pwo3* mutants using antibodies against H3K27me3 and H3. 10 days old, LD grown seedlings were analysed. Results are normalized to Col-0 and represent the mean of three biological replicates (error bars depict standard error) and asterisks indicate significantly different enrichment to Col-0 (p>0.05 for n.s.; p<0.05 for *, Student’s t-Test). See supplemental figure S4 for full dataset including control regions devoid of H3K27me3 and % IP values.

### The PWWP-domain of PWO1 is required for nuclear speckle formation in *N. benthamiana* and for interaction with H3

PWO1 contains a putative chromatin “reading” domain, the PWWP domain (Figure 1 and Supplemental Figure 1), which has been shown to target various proteins to chromatin (Dhayalan et al., 2010; Maltby et al., 2012; Wang et al., 2009), therefore this domain might have a similar function in the PWO1 protein. As PWO1 partially localizes to nuclear speckles (Figure 2F-G and I), we asked whether the PWWP domain may be required for subnuclear targeting. We therefore generated N-terminal (lacking the PWWP domain) and C-terminal deletions of PWO1 fused to GFP, which both contained the predicted NLS (*i35S_pro_::PWO1ΔPWWP-GFP* and *i35S_pro_::PWO1ΔC-GFP*), and compared subcellular localization with the full length PWO1 cDNA fused to GFP (*i35S_pro_::PWO1cDNA-GFP)* in transient expression assays in *N. benthamiana* (Figure 6A-C). Although fewer speckles were visible in C-terminally truncated PWO1 constructs (Figure 6B), only complete loss of speckle formation and partly cytoplasmic localization was observed with the PWO1 construct lacking the PWWP domain (Figure 6C).

**Figure 6:**
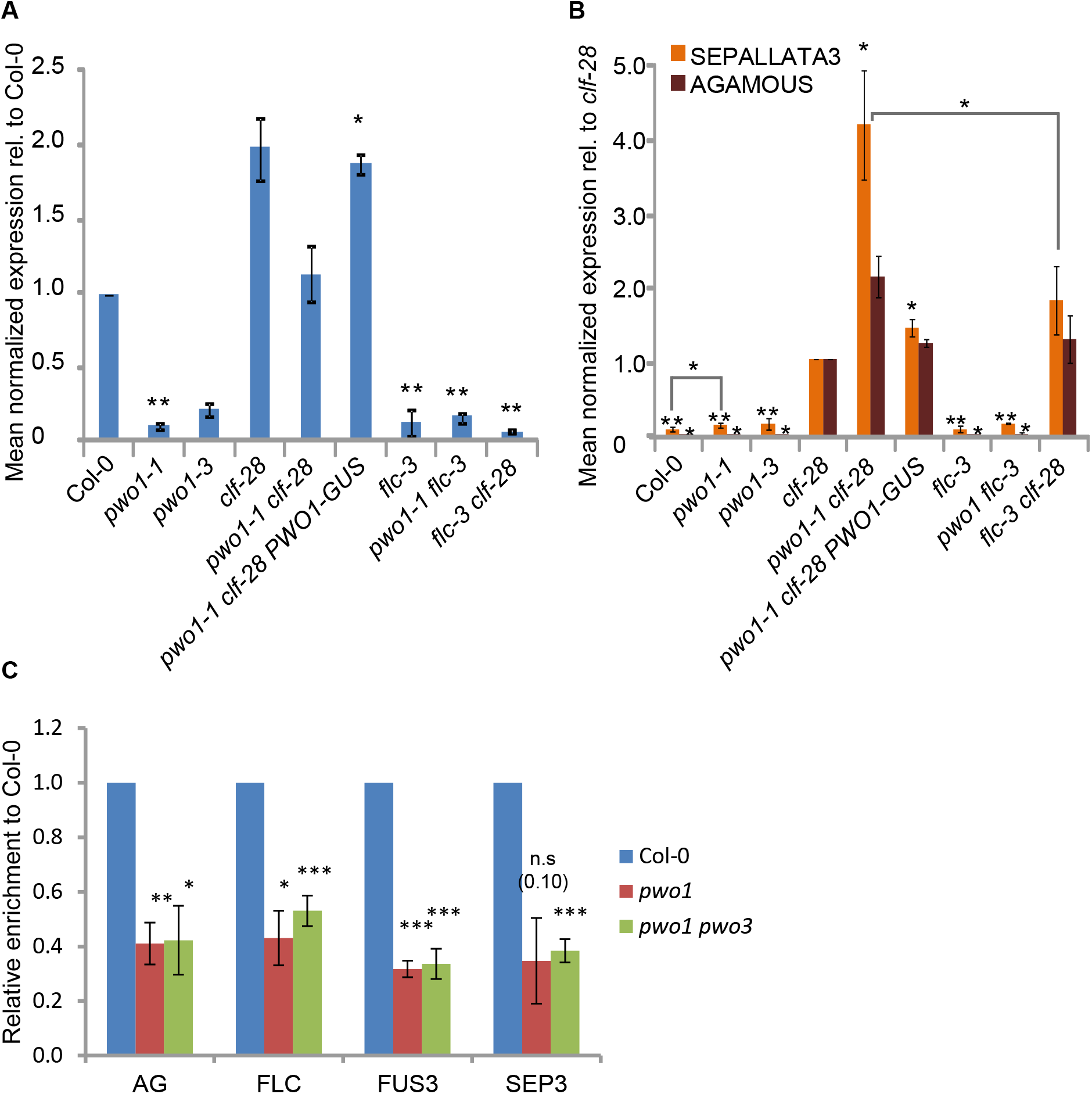
The PWWP domain of PWO1 is required for nuclear speckle formation and interaction with H3. **A - D.** Transient expression of PWO1-GFP variants in *N. benthamiana* leaf epidermal cells. **A.** *i35S_pro_:PWO1cDNA-GFP,* **B.** *i35S_pro_:PWO1 ΔC-GFP,* **C.** *i35S_pro_:PWO1 ΔPWWP-GFP.* Expression was induced with 5 μM β-estradiol for 5 hours. Scale bars: 20 μM. **D-F.** Co-IP of H3 with the full cDNA of PWO1 fused to GST (D), with a mutated version of PWO1 carrying a W63A point mutation (E) or the N-terminal part of PWO1 (F). 2% of input was run in the gel as loading control (I: input; IgG: beads coupled to IgG as negative control; beads: uncoupled beads). **G-I.** Comparison of *pwo1 pwo2 pwo3* mutants carrying different constructs: (G) *pwo1 pwo2 pwo3 PWO1::PWO1-GFP;* (H) left: *pwo1 pwo2 pwo3;* right: *pwo1 pwo2 pwo3 PWO1::PWO1_W63A_-GFP;* (I), *pwo1 pwo2 pwo3 PWO1::PWO1_W63A_-GFP* showing callus-like tissue in the SAM (asterisk) and leaf primordium with trichomes (arrow) which are never observed in *pwo1 pwo2 pwo3* triple mutants. G and H 2 weeks old plants, I 3 weeks old plant grown in tissue culture in long-day photoperiod. Scale bars: 1 mm.

As PWWP domains have been shown to confer binding to histones, for example in the mammalian *de novo* DNA methyltransferase DNMT3A (*Mus musculus)* or the Pdp1 protein from *Schizosaccharomyces pombe* (Dhayalan et al., 2010; Qiu et al., 2002; Wang et al., 2009) we next sought to determine whether PWO1-PWWP may also interact with histones. An alignment of the predicted PWO1 PWWP domain with those of the *Arabidopsis thaliana* proteins PWO2, PWO3 and ATX1, the mouse DNMT3A, DNMT3B and MSH6, the human NSD2 and the S. *pombe* Pdp1 proteins detected conservation of several important residues, which are predominantly hydrophobic (supplemental Figure S6). To further characterize the binding ability of the PWWP domain, the full PWO1 cDNA or a shorter fragment corresponding to the N-terminal half of the protein were fused to GST (Glutathione-S-transferase) and purified from *E. coli* to assay binding to H3 by co-immunoprecipitation (co-IP) experiments with anti-H3 antibodies. Both proteins were able to bind H3, indicating that the N-terminal part of the protein, which contains the PWWP domain, is sufficient to confer binding to H3 (Figure 6D and F). It has been shown that direct interaction of PWWP domains to histones occurs through a hydrophobic pocket formed by three aromatic amino acids (Qin and Min, 2014). PWO1-PWWP contains a partial hydrophobic pocket formed by two tryptophan residues (supplemental Figure S6). Strikingly, the point mutation of one of them, W63, to alanine (PWO1-W63A), decreases PWO1-H3 interaction *in vitro* (Figure 6E).

To analyze the role of the PWWP domain in PWO1’s functions the *pwo1 pwo2 pwo3* triple mutant was transformed with a construct carrying the point mutation in this domain (*PWO1::PWO1_W63A_-GFP*) and expression and nuclear localization of PWO1_W63A_-GFP was confirmed *in vivo* (supplemental Figure S7). The phenotypic analysis of two independent *PWO1::PWO1_W63A_-GFP pwo1 pwo2 pwo3* transgenic lines showed a partial complementation so that cotyledons and roots appeared more normal than in the triple mutants, i.e. hypocotyls and cotyledons were more elongated and cotyledons were accumulating chlorophyll. However, the shoot apical meristem (SAM) seemed initially reduced and then produced callus-like tissue from which leaves eventually developed (Figure 6 G-I).

Thus, the PWWP domain of PWO1 appears to be essential for its interaction to H3. In addition, PWO1-H3 interaction might be required for PWO1 nuclear localization and formation of nuclear speckles in *N. benthamiana* and for full PWO1 activity in Arabidopsis.

### The PWO1 PWWP domain binds to histone peptides lacking H3S28 phosphorylation

In order to analyze whether the PWO1 PWWP domain confers binding to posttranslationally modified histones, we fused it to GST (Glutathione-S-transferase), expressed it in *E. coli* and purified the fusion protein. The GST-PWO1-PWWP fusion protein was then incubated with the MODified^TM^ histone peptide array harboring different combinations of naturally occurring histone modifications on histone peptides (Bock *et al.,* 2011). The GST-PWO1-PWWP fusion protein bound rather non-specifically to histone peptides and modified histone peptides, however, binding was specifically inhibited by phosphorylation of H3S28 (Figure 7A). Importantly, nonhistone peptides were not bound and binding was not disrupted by H3S10 phosphorylation, which lies in a similar sequence context as H3S28 (N-ARKS-C). This suggests that loss of binding is not only caused by an increase in negative charge, but also depends on the sequence context N- or C-terminal to the ARKS sequence.

**Figure 7:**
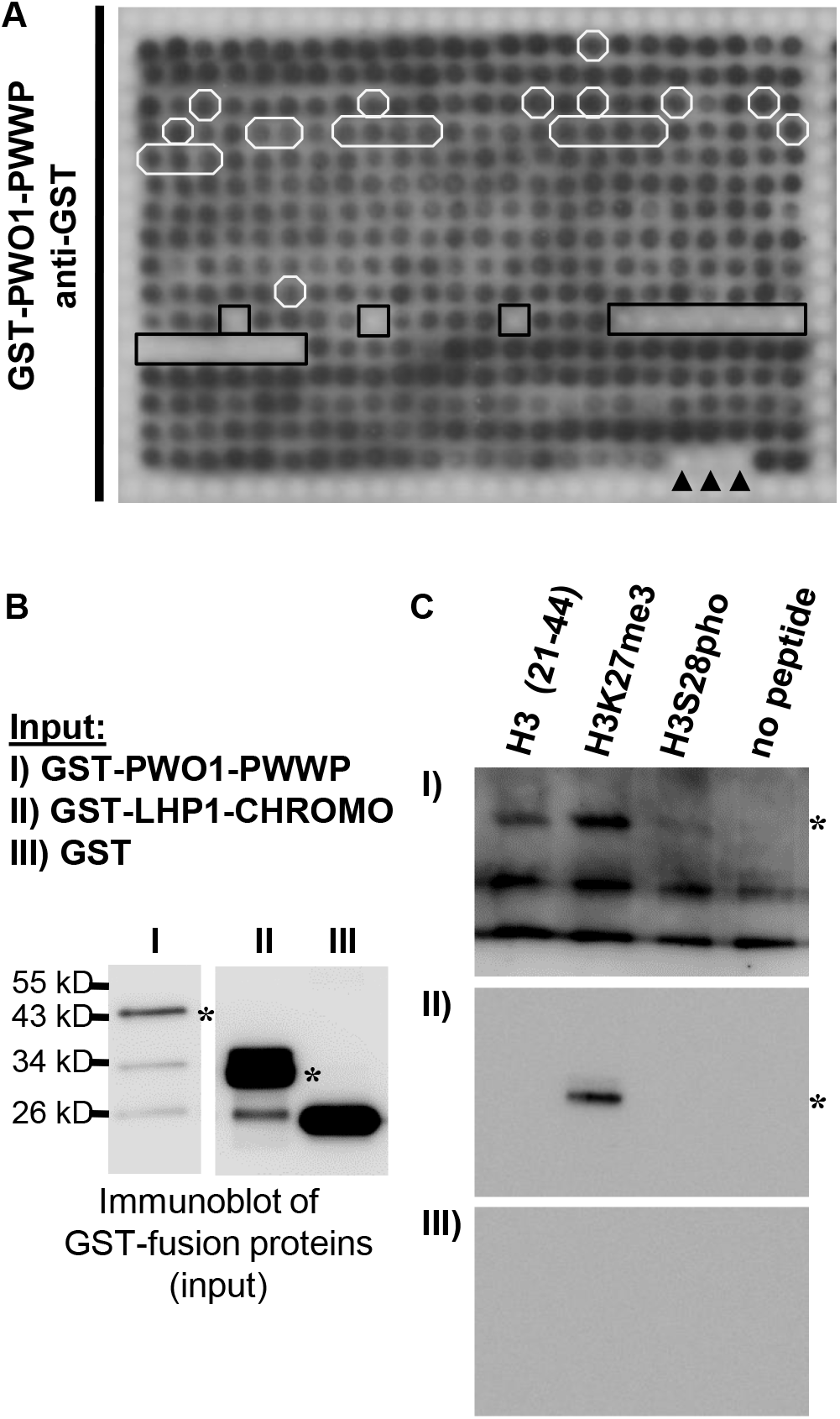
The PWO1 PWWP domain binds to histone peptides lacking H3S28 phosphorylation. **A.** Binding of the PWO1 PWWP domain (GST-PWO1-PWWP) to peptide arrays containing 384 different peptides. Peptides containing H3S10p are in white circles, peptides carrying H3S28p are in black boxes. Arrowheads indicate non-histone peptides. For a detailed annotation of all spots cf. http://www.activemotif.com/catalog/668/modified-histone-peptide-array. **B., C.**, Confirmation of PWO1-PWWP histone binding by peptide pulldown analyses. GST-fusion proteins were separated by polyacrylamide gel electrophoresis and blotted to a Nitrocellulose membrane (B). Fusion proteins were detected with anti-GST antibody staining. **C**, I. –III fusion proteins were incubated with biotinylated histone peptides and precipitated with streptavidine beads. Pulled down proteins were separated by polyacrylamide gel electrophoresis, blotted to a Nitrocellulose membrane and detected with anti-GST antibody staining, asterisks indicate fusion proteins in B and C. Faster migrating bands in I) are GST-PWO1 degradation products.

The binding capacity of GST-PWO1-PWWP to unmodified and modified histone H3 peptides was then independently demonstrated *in vitro* in peptide pull-down experiments. As revealed in the peptide array, H3 (aa 21 - 44) and H3K27me3 peptides precipitated the GST-PWO1-PWWP fusion protein, however, binding was strongly reduced by H3S28 phosphorylation (Figure 7B-C). While GST alone was not able to bind to the biotinylated peptides, a GST fusion protein with the CHROMO domain of TFL2/LHP) (GST-TFL2-CHROMO), showed *in vitro* binding ability to H3 peptides trimethylated at K27 consistent with previously published data (Turck *et al.,* 2007; Zhang *et al.,* 2007b).

Although currently the impact of H3S28 phosphorylation on PcG mediated H3K27me3 and gene silencing has not been studied in plants, a role for H3S28p in antagonizing PcG silencing has recently been uncovered in animals (Gehani *et al.,* 2010; Lau and Cheung, 2011). As H3S28 is adjacent to the PRC2 modified H3K27 also in plants, a similar mechanism of PcG displacement may occur here as well.

## Discussion

### PWO1 may recruit Arabidopsis PcG proteins to subnuclear speckles

Plant genomes contain a multitude of genes with similarity to chromatin regulators. However, only a few plant-specific components of chromatin and PcG complexes have been identified so far. These may substitute for non-conserved proteins and contribute to key processes in epigenetic gene regulation including recruitment of the complexes to their target genes and inheritance of the epigenetic state through mitosis and/or meiosis. Alternatively, they may fulfill plant-specific roles which may have evolved to accommodate the different lifecycles of plants and animals.

In this study, we aimed to identify interaction partners of the PRC2 protein CLF and focused on plant-specific components with putative chromatin “reading” domains. Among other proteins, we identified the PWWP domain protein PWO1 which interacts with all three histone methyltransferase subunits of the Arabidopsis PRC2, CLF, SWN and MEA in yeast. PWWP domains belong to the “Royal” family of domains which have diverse functions in chromatin regulation and have been shown to bind to DNA or (post-translationally modified) histones (Dhayalan et al., 2010; Maurer-Stroh et al., 2003; Qin and Min, 2014; Qiu et al., 2002; Wang et al., 2009). PWWP domains are found in a diverse range of proteins in uni- and multicellular organisms including Arabidopsis whose genome encodes for at least 19 PWWP domain proteins (16 identified in (Alvarez-Venegas and Avramova, 2012) plus the three in this study). Similar to PcG proteins, *PWO1* is expressed in diverse tissues and localizes to euchromatic regions in the nucleus. Interestingly, we revealed that PWO1 localizes to nuclear speckles when expressed transiently in *N. benthamiana* and stably in Arabidopsis, however, the distribution of the speckles in Arabidopsis seemed more uniform despite their absence from chromocenters. CLF and PWO1 influence each other’s subnuclear localization in tobacco: while CLF only localizes to nuclear speckles in the presence of PWO1, presence of PWO1 in larger nuclear patches is only observed when co-expressed with CLF. Nuclear speckle formation depends on the PWO1 PWWP domain suggesting that chromatin targeting by the PWWP domain may precede speckle formation. The identity of the speckles is currently unclear (Del Prete et al., 2015), but also other PcG proteins like VRN2, LHP1/TFL2 and EMBRYONIC FLOWER 1 (EMF1) were found to localize to nuclear speckles when expressed transiently in *N. benthamina, N. tabacum* or onion epidermal cells (Calonje et al., 2008; Gaudin et al., 2001; Gendall et al., 2001; Libault et al., 2005; Zemach et al., 2006). Nevertheless, as the speckles are not clearly detected in Arabidopsis, we cannot exclude that the transient expression of PWO1 in a heterologous system may favor speckle formation. It is also possible that putative PWO1 speckles in Arabidopsis are much smaller than the ones detected in *N. benthamiana* which will require higher resolution microscopy. Nevertheless, *Drosophila* and mammalian PRC1 proteins are found in so-called PcG bodies, which are observed as nuclear speckles whose number and size varies depending on the cell type (Pirrotta and Li, 2012). While the mammalian orthologue of CLF, ENHANCER OF ZESTE HOMOLOG2 (EZH2) is required for PcG body formation (Hernandez-Munoz *et al.,* 2005), EZH2 is not found in the bodies (Sewalt et al., 1998). However, there are still no clear evidences for PcG bodies in plants and further experiments will be required to determine whether PWO1-CLF speckles correspond with clustering of both proteins and interaction with PcG target genes as observed in Drosophila (Bantignies et al., 2011). We also found that PWO1 may form homodimers which could contribute to subnuclear speckle formation and possibly polymerization as shown for mammalian polyhomeotic homologue 2 (PHC2) (Isono et al., 2013).

### Role of the PWO family in Arabidopsis development, flowering time control and PcG target gene regulation

Plant PcG proteins control various developmental processes including flowering time, embryo and seedling development (Mozgova et al., 2015). Similar to *clf* and *emf* mutants, *pwo1* shows an early flowering phenotype in which mis-expression of some PcG target genes seem to play an important role. In contrast to *clf* and other PRC2 mutants, *FLC* expression is lower in *pwo1* (Jiang et al., 2008; Lopez-Vernaza et al., 2012), while other flower-specific PcG target genes are ectopically expressed in *pwo1* or show enhanced ectopic expression in *clf pwo1* mutants. Similar expression patterns which include reduced levels of *FLC* are also observed in *incurvata2, blister, chr11/17* and *ringlet* mutants (Barrero et al., 2007; Li et al., 2012; Schatlowski et al., 2010) whose wild-type products all show a physical and/or genetic interaction with PcG proteins or mutants, respectively. Thus, these and the PWO1 proteins may have a dual function as activators and repressors of distinct PcG target genes. Alternatively, the effect on *FLC* may be indirect and these proteins may repress a repressor of *FLC,* similar to what has been reported for the PcG target gene *FT* whose expression is down-regulated in strong PcG mutants (Farrona et al., 2011). Our chromatin analyses suggest that the observed reduction in H3K27me3 occupancy in *pwo1* mutants is largely due to a reduction in H3 occupancy, suggesting that *PWO1* is not required for H3K27me3 activity of PRC2. However, it may be involved in compaction of Pc-G target chromatin, either to facilitate PRC2 H3K27 methyltransferase activity or to compact nucleosomes after the H3K27me3 mark has been set. PWO1’s general affinity for histones is consistent with this function and it will be important to study interactions of PWO1 with chromatin remodeling components. In addition, it remains to be elucidated whether the decrease in H3 occupancy occurs throughout the genome or specifically at Pc-G target genes.

While *pwo1* mutants are similar in size to wild-type, they strongly enhance *clf* mutants resulting in small plants with severe leaf curling and very early flowering. The full function of PWO1 in plant development is, however, masked by the redundancy with its two homologous proteins PWO2 and PWO3. The triple mutants show shoot and root meristem arrest and produce no or severely affected post-embryonic organs and die a few weeks after germination indicating their essential role for development. Strong PcG mutants like *clf swn* double show a somewhat weaker phenotype as they keep on proliferating after germination and produce leaf and root tissue (Chanvivattana et al., 2004). This difference in phenotype may be explained by different PcG target gene activation in *pwo1/2/3* and PRC2 mutants or a PcG independent role of the PWO family. Nevertheless, the severe *pwo1/2/3* mutant phenotype reveals an essential role for the PWO family in preventing premature differentiation and maintaining meristematic activity.

### PWO1 interacts with H3 through its conserved PWWP domain and may recruit PcG proteins to unphosphorylated H3S28 tails

PWWP domains are found in numerous proteins and belong to the “Royal Family” of chromatin readers (Maurer-Stroh et al., 2003). Several PWWP domain proteins bind to histones, like the PWWP domain of DNMT3A which is able to interact with H3 or S. *pombe* Pdp1’s PWWP domain which binds H4, while others interact with DNA, such as DNMT3B (Dhayalan et al., 2010; Qiu et al., 2002; Wang et al., 2009).

We revealed that PWO1 is able to interact with H3 *in vitro* and that a fragment of the protein containing the PWWP domain is sufficient to allow this interaction. Binding of PWWP domains to histones occurs through a hydrophobic pocket formed by three aromatic residues which in many cases have been shown to specifically recognize methyl-lysine residues on the histone tail (Qin and Min, 2014). Sequence analysis of the PWWP in PWO1-3 showed that the domain in these proteins have a partial hydrophobic domain in which the first aromatic residue has been substituted by glycine. Other proteins with an incomplete aromatic cage, such as the Retinoblastoma-Binding Protein 1 (RBBP1), do not show specificity for methylated residues (Gong et al., 2012), indicating that a different mechanisms might be also involved in PWO1 -H3 binding.

Further demonstration of the importance of the PWWP domain of PWO1 with H3 was shown through the point mutation of the W63 residue of PWO1’s semi-aromatic cage. This amino acid has been described to form part of the groove that interacts with specific residues in the N-terminal tail of H3 (Qin and Min, 2014). When this residue was mutated to alanine in PWO1-PWWP a decrease of the binding of the domain to H3 was observed *in vitro.* Mutation of the corresponding residue in the Pdp1 protein of S. *pombe* strongly disrupted *in vitro* binding of Pdp1 to H4K20me3 (Wang et al., 2009). Therefore, our data indicate that other residues in PWO1’s PWWP may also play an important role in the interaction with H3 under the tested conditions. This hypothesis was also supported by our *in vivo* results for complementation of the *pwo1 pwo2 pwo3* mutants with the version of PWO1 carrying the W63A point mutation. The experiments showed that the PWO1-W63A was not able to fully complement the triple mutants, probably indicating a partial activity of the mutated PWWP domain and of the protein carrying this mutation. This residue might be especially important for the function of PWO1 in maintaining meristematic activity, considering that the SAM was the organ most strongly affected in the *PWO1::PWO1_W63A_-GFP pwo1 pwo2 pwo3* plants.

We further revealed, that at least *in vitro,* PWO1 does not have a preference for posttranslationally modified histone peptides, however, the interaction is inhibited by phosphorylation of H3S28. A related function has been shown for a cysteine-rich domain of DNMT3L which specifically recognizes non-methylated H3K4 (Ooi *et al.,* 2007).

Currently there is limited information available on the function of H3S28p or H3K27me3S28p in plants. H3S28p as well as H3S10p accumulate during mitosis and meiosis in diverse plant species, but are hardly detectable in interphase nuclei (Gernand *et al.,* 2003), similar to what has been observed in mammals (Goto *et al.,* 1999). Importantly, H3K27me3S28p was shown to counteract PcG repression upon stress induction (Gehani *et al.,* 2010; Lau *et al.,* 2011), potentially providing a powerful way of quickly and transiently relieving PcG repression. PWO1 may therefore be involved in stabilizing the PRC2-Histone interaction which can be disrupted by phosphorylation of H3S28.

In conclusion, PWO1 (and possibly also PWO2 and PWO3) may serve as factors which interact with PcG proteins to recruit them to subnuclear speckles and to mediate full H3/nucleosomal compaction. In this activity its PWWP domain may act as key element in the interaction of PWO1 with the chromatin, possibly by its ability to interact with histones. Accumulation of H3S28p during stress treatment and/or cell division may lead to displacement of PWO1 and associated PcG proteins and release of PcG target gene silencing. The *PWO1/2/3* genes are essential for proper cell division and maintenance of stem cells as lack of their function causes early seedling lethality with root and shoot meristem arrest. Whether this is a result of improper recruitment of PcG proteins is an interesting possibility to explore in the future.

## Methods

### Biological material

The *PWO1* T-DNA insertion lines were identified using the SIGNAL database (http://signal.salk.edu/cgi-bin/tdnaexpress) and provided by the Nottingham Arabidopsis Stock Centre: *pwo1-1* (N815951), *pwo1-2* (N420954), *pwo2-2* (N636093), *pwo3-2* (N836957) (Alonso et al., 2003; Sessions et al., 2002). TILLING (Targeting Induced Local Lesions IN Genomes) mutants were ordered from the Seattle Tilling Project (STP) and analyzed (http://tilling.fhcrc.org) (Till et al., 2003). The *pwo1-3* (N93526) allele exhibits a point mutation (R46Stop) leading to a premature stop codon. Homozygous mutants were isolated by PCR-based genotyping (for oligonucleotide sequences, see supplemental Table S1). For analysis of genetic interactions crosses were performed with *clf-28* (N639371) and *flc-3* (Michaels and Amasino, 1999). All genotypes used in this study are in the Columbia-0 (Col-0) background.

Seeds were sterilized and sown on GM media (half-strength Murashige and Skoog medium plus 0.5% sucrose), stratified for 2 d at 4°C and transferred to soil after 10 to 12 d. Plants were grown at either long-day conditions (16 hlight/8 h dark cycles at 20°C) or short-day conditions (8 h light/16 h dark cycles at 20°C).

Transformation of *Arabidopsis thaliana* was performed after the floral dip method using the *Agrobacterium tumefaciens* strain GV3101 pMP90 (Clough and Bent, 1998).

### Complementation of the *pwo1-1* and *pwo1 pwo2 pwo3* mutant phenotype

The *PWO1_pro_:PWO1-GUS* and *PWO1_pro_:PWO1-GFP* constructs where generated by amplification of the genomic locus of *PWO1* including 1534 bp upstream of the transcriptional start site using the following primers: CTAACTTCACAGCACGGCTCTGAGG and TTGAACTCTTCTTCTCTCGTTAAAGGC. The PCR fragment was cloned into the pCR8/GW/TOPO entry vector (Invitrogen) and subsequently cloned into pMDC163 as translational fusion with the uidA gene, and into pMDC107 as translational fusion with GFP (Curtis and Grossniklaus, 2003). The *PWO1_pro_:PWO1_W63A_-GFP* construct was created using the *PWO1_pro_:PWO1-GFP* cloned in the pCR8/GW/TOPO entry vector and using primers GTGCTTAGGGATGCGTACAATTTAGAG and ATTAAAATACGAATGCTTTCAGTAATC to introduce the mutation.

Using the floral dip method, *pwo1-1^-/-^ pwo2-2^-/-^ pwo3-2-/-* plants were transformed with *A. tumefaciens* carrying the T-DNA vectors. Plants carrying the transgene were selected on GM medium containing hygromycin (15 mg/ml) and further segregated to obtain the desired genotype.

### Phenotypic Analyses and Imaging

Photographs were taken with an AxioCam ICC1 camera (Zeiss) mounted onto a Zeiss Stemi 2000C. For scanning electron microscopy, plant material was treated as described previously (Kwiatkowska, 2004) and SEM was performed using the LEO (Zeiss) microscope and software. For visualisation of chromocenters in *A. thaliana*’s PWO1 ::PWO1-GFP and PWO1::PWO1(S63A)-GFP lines, anthers’ filaments were mildly fixed in 4% PFA under vacuum for 2min, washes 3 times with PBS, incubated with propidium iodide solution (1ug/ml) for 20min in darkness and again washed 3 times with PBS. Fluorescence was monitored with a Zeiss LSM 710 confocal laser scanning microscope. Intensity values of fluorescence in particular regions of nuclei were scored afterwards using Plot Profile feature for intensity profiles in Fiji/ImageJ 1.48p.

For flowering time analysis genotypes were grown in parallel under indicated conditions and rosette leaf number before bolting was analysed for at least 15 plants per genotype.

### Yeast Two-Hybrid Interaction Studies

For interaction studies of PWO1 with CLF, SWN and MEA, the vectors pGAD-PWO1-CDS, pGBD-PWO1-CDS and pGAD-SWNΔSET were generated. Additional vectors used were pBD-CLFΔSET (Chanvivattana et al., 2004) and pAD-MEA-CDS (Lindner et al., 2013).

To generate pGAD-PWO1-CDS, pGBD-PWO1-CDS, a full-length cDNA was ordered (Riken RAFL16-55-O22) and used as template for PCR amplification of the complete PWO1 coding sequence (For: ATGGCAAGTCCAGGATCAGGTGC, Rev:TTGAACTCTTCTTCTCTCGTTAAAGGC) which was cloned into pCR8/GW/TOPO entry vector and subsequently recombined into the vectors pGBKT7-DEST and pGADT7-DEST (Horak et al., 2008).

All yeast techniques were performed as described in the ‘Yeast Protocols Handbook’ (Clontech Laboratories, Inc., protocol PT3024-1, version PR13103). The yeast strains YST1 and AH109 (were transformed with Gal4-BD and Gal4-AD constructs. After mating, yeast two-hybrid studies were performed by dilution series on selective media.

### Co-immunoprecipitation assays

4 g of 2-week-old Col-0, *PW01_pro_:PWO 1-GFP* and *PWO1_pro_.PWO1-GFP 35S_pro_:CLF-mCherry* seedlings were harvested and nuclear proteins were extracted from samples as in (Smaczniak et al., 2012). To immunoprecipitate GFP fused proteins, nuclear proteins were incubated for 2 h at 4°C with 50 μl of μMACS anti-GFP MicroBeads (Miltenyi Biotec). Beads were immobilized in calibrated μcolumns (Miltenyi Biotec), washed 6x with 200 μl of Lysis Buffer and 2x with 200 μl of Wash Buffer 2 and, finally, bound proteins were eluted from the μcolumns with Elution Buffer pre-heated at 95°C (μMACS GFP Isolation Kit; Miltenyi Biotec). Eluted proteins were run on a 10% SDS-PAGE gel and transferred to PVDF membranes. Membranes were incubated with anti-DsRed antibody and anti-rabbit IgG coupled to HRP (Supplemental Table S3) as primary and secondary antibodies, respectively. SuperSignal West Femto Chemiluminiscent Substrate (Thermo Scientific) was used to develop the membranes and the signal was analyzed in a Fuji ImageQuant LAS4000.

*PWO1-CDS* and *PWO1_W63A_-CDS* constructs were cloned in the pGEX-4T-3 vector (GE Helthcare) modified to be used with the Gateway system (Invitrogen). The plasmids were expressed in *E. coli* BL21 strain and proteins were purified using Glutathione-Sepharose 4B (Sigma, #GE17-0756-01). Anti-H3 antibodies (Millipore, #05-928) were coupled to Dynabeads® Protein A (*life* technologies™, #10001D) and subsequently incubated with 5-7 ng of PWO1-GST or PWO1_W63A_-GST proteins in 1x PBX during 4 hours at 4°C. After incubation, beads were washed three times with 1x PBS and resuspended in 2x SDS-PAGE loading buffer. Proteins were loaded in 10% SDS-PAGE gels and transferred to PVDF membrane. Membranes were developed with anti-GST (Sigma, #G7781) and anti-H3 (Diagenode, #C15200011) antibodies.

### Transient Colocalization Assay

Modified versions of pMDC7 carrying the GFP (pABindGFP) or mCherry (pABindCherry) coding sequence (Bleckmann et al., 2010) were used to insert the complete coding sequence of *PWO1* as well as truncations of *PWO1, CLF* and *SWN* via Gateway-cloning (Invitrogen).

Vectors were transformed in *A. tumefaciens* GV3101 pMP90 carrying the silencing suppressor p19. For transient expression assays, abaxial sides of leaves of 4-week-old *Nicotiana benthamiana* plants were infiltrated with *A. tumefaciens* suspension as described by (Bleckmann et al., 2010). 48 to 72 h after *A. tumefaciens* infiltration, expression was induced by brushing 20 mM β-estradiol in 0.1% Tween onto infiltrated leaves. To limit overexpression artefacts, fluorescence was monitored in leaf epidermis cells after a short induction period (4 to 6 h when fluorescence was visible) using a Zeiss LSM 710 confocal laser scanning microscope. Induction times for testing localization of different PWO1 variants were kept similar (fluorescence monitoring after 5 h of induction).

### Gene Expression Analyses

Detection of GUS activity was performed as described previously (Colon-Carmona et al., 1999).

Pools of 10 days old, LD grown seedlings were used for total RNA extraction (RNeasy plant mini kit; Qiagen). RNA was resuspended in 50 μL RNase-free water, treated with DNase (Fermentas), transcribed into cDNA using SuperScriptIII following the manufacturer’s instructions (Invitrogen), and subjected to real-time PCR. qRT-PCR analyses was performed with technical triplicates and three biological replicates (grown and harvested independently) using oligonucleotides listed in Supplemental Table S2. The Mesa Blue Sybr Mix (Eurogentec) was used for amplification in a Chromo4 realtime PCR cycler (Biorad). Expression levels were normalized to the reference gene At4g34270 (nblack) (Czechowski et al., 2005).

### Chromatin Immunoprecipitation

Chromatin immunoprecipitations from 2g of 2-week-old Col-0, *pwo1-1* and *pwo1-1/3-* 2 pools of 10 days old, LD grown seedlings were performed using Plant ChIP kit (Diagenode #C01010150), following instructions given in the kit’s protocol. Shearing of chromatin was obtained by sonicator Bioruptor Pico (Diagenode, #B01060001). For immunoprecipitations anti-H3K27me3- and anti-H3 specific antibodies were employed (see supplemental Table S3). Three biological replicates were grown and harvested independently.

qPCR analyses were done with SYBR Green I Master mix (Roche, #04887352001) and iQ5 cycler detection system (Biorad, #170-9780), using a two-step program. Ct values from input and immunoprecipiated samples were obtained in 3 technical replicates. Differences between genotypes were scored by comparison of % of input and standard error values. Sequences of oligonucleotides used for ChIP analyses are listed in Supplemental Table S2.

### Immunofluorescence

Roots of ten days old seedlings grown on GM medium were harvested, fixed and nuclei were isolated essentially as described in (Lysak et al., 2006). PWO1-GFP nuclear distribution was analysed using immunofluorescence with anti-GFP antibodies and secondary antibodies as depicted in Supplemental Table S3 according to (Lysak et al., 2006). DAPI-staining of the nuclei was performed as described in (Lysak et al., 2006).

### Precipitation of GST-fusion proteins with biotinylated peptides

Protein domains were expressed as GST-fusion proteins (backbone pGEX4T3) in *E. coli* strains Rosetta(DE3). GST-fusion protein expression was induced with 0.5 mM IPTG for 3 hours at 28°C. Proteins were purified using the MagneGST Pulldown system (Promega) according to the manufacturer’s instructions. Purified GST-fusion proteins were used for precipitation experiments with biotinylated histone peptides listed in Supplemental Table S3 as in (Shi *et al.,* 2006) with modifications: 1 μg of fusion protein was incubated with 1 μg of biotinylated peptide in 300 μl Gozani-buffer (50mM Tris-HCl pH7,5; 300mM NaCl; 0,1% Igepal; 1mM PEFA and 1:100 plant protease inhibitor cocktail (Sigma-Aldrich P9599)) over night at 4°C on a rotating platform. Per sample 15 μl of Streptavidin magnetic beads (Invitrogen) were added and incubated for 1 h at 4°C on a rotating platform. Beads were washed 3 times with 500 μl Gozani-buffer at 4°C, then the bound proteins were eluted with 50 μl 0,1% SDS and incubation at 95°C for 5 min. The eluted proteins were studied by immunoblot analysis.

### ModifiedTM Histone Peptide Array

The purified GST-PWO1-PWWP fusion protein was hybridized to a ModifiedTM Histone Peptide Array (Active Motif) to reveal binding specificity to modified histone peptides. The array, hybridization and analysis were performed as described in (Bock *et al.,* 2011).

### Antibodies and peptides

Antibodies and peptides used in this study are listed in supplemental Table S3.

### Accession numbers

Sequence data from this article can be found in EMBL/GenBank data libraries under the accession numbers at3g03140 (PWO1), at1g51745 (PWO2), at3g21295 (PWO3), at5g10140 (FLC), at2g23380 (CLF), at4g02020 (SWN), at1g02580 (MEA), at5g17690 (TFL2/LHP1), at1g24260 (SEP3), at4g18960 (AG), at1g65480 (FT).

## Acknowledgements

We gratefully acknowledge funding to support this research by the Deutsche Forschungsgemeinschaft through grant SFB973, the Boehringer Ingelheim Foundation and the European Commission Seventh Framework-People-2012-ITN project EpiTRAITS (Epigenetic regulation of economically important plant traits, no-316965). We thank Justin Goodrich for support for the initial yeast two-hybrid screen, Nora Lorberg for excellent technical assistance, José Luis Riechmann for providing the yeast two-hybrid cDNA library, Andrea Bleckmann for the modified pMDC7 vectors and the Nottingham Arabidopsis Stock Centre for seeds and ABRC for DNA stocks. We are further grateful to Celine Sabatel and Jean-Jacques Goval of Diagenode for their support and attention to protocol development for low chromatin amounts. We thank Justin Goodrich and members of the Schubert lab for critical reading of the manuscript.

## Author contributions

M.L.H., P.W., S.F. and D.S. designed the research, M.L.H., S.F., P.M., O.K., C.K. and I.K. performed the experiments, I.K. and A.J. provided reagents, M.L.H., S.F. and D.S. wrote the paper.

## Supplemental data

The following materials are available in the online version of this article:

Supplemental Figure S1: Alignment of PWO1, PWO2 and PWO3 proteins

Supplemental Figure S2: *PWO1, PWO2* and *PWO3* gene structures, mutant alleles and phenotypes of single and double mutants and complemented lines

Supplemental Figure S3: Analysis of genetic interaction of *pwo1* and *clf* in SD conditions

Supplemental Figure S4: Analysis of *FT* expression

Supplemental Figure S5: H3 and H3K27me3 analyses in *pwo* mutants

Supplemental Figure S6: Alignment of PWWP domains of Arabidopsis, mouse, human and *S. pombe* proteins

Supplemental Figure S7: PWO1-GFP signal intensity is not affected by the mutation of its PWWP domain

Supplemental Table S1: Oligonucleotides for genotyping Supplemental Table S2: Oligonucleotides for RT- and ChIP-PCR Supplemental Table S3: Antibodies and peptides used in this study

